# Hierarchical control of microbial community assembly

**DOI:** 10.1101/2021.06.22.449372

**Authors:** Sammy Pontrelli, Rachel Szabo, Shaul Pollak, Julia Schwartzman, Daniela Ledezma-Tejeida, Otto X. Cordero, Uwe Sauer

## Abstract

Metabolic processes that fuel the growth of heterotrophic microbial communities are initiated by specialized biopolymer degraders that decompose complex forms of organic matter. It is unclear, however, to what extent degraders control the downstream assembly of the community that follows polymer breakdown. Investigating a model marine microbial community that degrades chitin, we show that chitinases secreted by different degraders produce oligomers of specific chain lengths that not only select for specialized consumers but also influence the metabolites secreted by these consumers into a shared resource pool. Each species participating in the breakdown cascade exhibits unique hierarchical preferences for substrates, which underlies the sequential colonization of metabolically distinct groups as resource availability changes over time. By identifying the metabolic underpinnings of microbial community assembly, we reveal a hierarchical crossfeeding structure that allows biopolymer degraders to shape the dynamics of community assembly.

**One sentence summary:** Specialized biopolymer degraders direct the trajectory of microbial community assembly through interconnected modes of nutrient crossfeeding.

## Introduction

Microbial communities mediate a staggering number of biological processes that contribute to the health of humans, animals(*1*) and the planet as a whole(*2*). The metabolic activity of co-occurring species results in the formation and depletion of nutrients in shared resource pools, where different modes of competition and cooperation impact community composition and form a network of metabolic crossfeeding(*3–24*). Individual species often provide specific metabolic functions that influence the growth of a broader community, including use of terminal electron acceptors(*6, 25, 26*), degradation of complex carbon sources(*27–30*), removal of toxic byproducts(*31–33*), or assimilation of nitrogen and sulfur(*10, 34–36*). However, we remain with a limited molecular understanding of the extent to which individual species impact the formation and structure of crossfeeding networks and community assembly. Here, we demonstrate multiple metabolic mechanisms by which specialized biopolymer degraders influence the trajectory of community assembly, exemplified with an 18 member community that thrives on chitin, the second most abundant polysaccharide on the planet(*37*). Similar to other polymer degrading communities(*27–29*), chitin communities require specialized degraders to supply nutrients to nonspecialized downstream consumers. Our community consists of phylogenetically diverse seawater isolates with distinct metabolic capabilities, which become abundant at different points during assembly of chitin communities. This provides a system that allows us to demonstrate how specialized chitin degraders initiate a hierarchal food web that shapes the community during the assembly process.

### Functional classification of bacterial isolates

We first classified the 18 isolates, collected from natural polymer degrading communities(*15, 27*)(Table S1), into three functional guilds based on their phenotypic capabilities for chitin utilization (Fig 1A). These species were tested for the ability to grow on chitin, the chitin monomer *N*-acetylglucosamine (GlcNAc), and the dimer chitobiose (Fig 1AB, S1, S2): Five degraders used all three substrates as a sole carbon and nitrogen source, and six exploiters could not use chitin but grew on GlcNAc, four of which also grew on chitobiose. Seven scavengers could not use any of these substrates; hence, they must rely on nutrients secreted by other species to thrive in the community. Previously published genome data of these 18 isolates(*15, 27*) revealed a wide range of chitin utilization potential (Fig 1A) that closely matches phenotypes. All five degraders, and two exploiters contain chitinases, uptake components specific for GlcNAc, and all genes required for conversion of GlcNAc into glycolytic intermediates: GlcNAc kinase, GlcNAc-6-phosphate deacetylase, and glucosamine-6-phosphate deaminase. The remaining four exploiters contain all metabolic components required to metabolize and transport GlcNAc (Fig 1A). All seven scavengers lack at least one transporter or catabolic gene required for GlcNAc utilization (Fig 1A). These varying metabolic capabilities prime guilds to become abundant at different points during community succession due to the different nutrients that become available over time(*15*). To visualize the trajectory of each functional guild during community assembly, we mapped 16s sequences of all species to 16s ribosomal gene exact sequence variants whose abundance changes were previously determined on chitin particles colonized from a large species pool in seawater(*27*). The mean trajectories of each functional guild revealed unique dynamics (Fig 1C), whereby degraders are the first to reach their maximum abundance, followed by exploiters and finally scavengers. This succession pattern portrays an order in a trophic cascade; i. e. degraders cause the release of chitin degradation products that are consumed by exploiters, which in turn release broader sets of metabolites that support scavengers.

**Figure 1:**
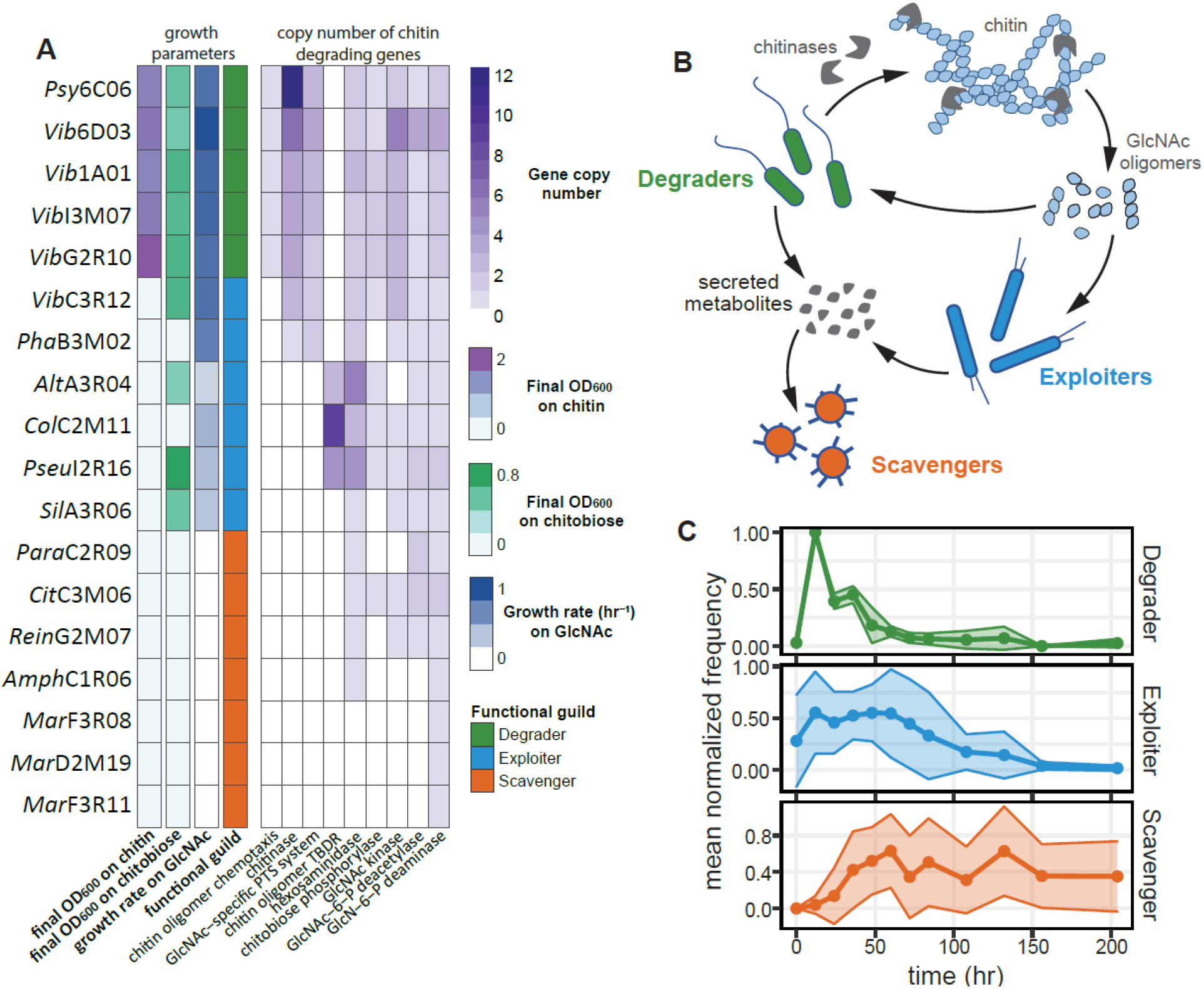
Successional dynamics of chitin degrading seawater communities. A) Species used in this study. Heatmap depicting (left to right) final OD_600_ after growth on 2 g/l colloidal chitin, final OD_600_ after growth on 10 mM chitobiose, growth rate on 20 mM GlcNAc, classified functional guild, and copy numbers of genes relevant for chitin degradation. Growth data was obtained from three independent biological replicates. B) Schematic of functional guilds: Degraders can grow on chitin as a sole carbon source through breakdown of chitin using chitin degrading enzymes, exploiters can grow on monomeric or oligomeric N-acetylglucosamine (GlcNAc) as a sole carbon source, and scavengers require metabolites secreted by other species to be sustained in the community. C) 16s sequences of the 18 investigated species are mapped to 16s ribosomal gene exact sequence variants whose abundances were previously determined on chitin particles colonized from a species pool in seawater(*27*). Mean normalized frequencies of species that comprise each functional guild are plotted over time. TBDR, TolB-Dependent receptor

### Downstream metabolic influence of secreted chitinases

It is likely that degraders form publicly available products that can support the growth of exploiters and possibly select for specific species. To test this, extracellular enzymes were concentrated from culture supernatants of each degrader during growth on chitin and used to digest fresh colloidal chitin. Liquid chromatography mass spectrometry (LC-MS) quantification of oligosaccharides with six or fewer residues revealed that each digest resulted in a unique oligomer profile, with the disaccharide chitobiose and the monomer GlcNAc as the most abundant products (Fig 2A). Each of the six exploiters was then grown on the five chitin digests (Fig 2B, Fig S3). *Sil*A3R06 and three other exploiters grew well on all digests. While *Pha*B3M02 and *Alt*A3R04 hardly grew on the *Psy*6C06 digest that contained chitobiose as the major oligomer (Fig 2A), but grew on *Vib*1A01 and *Psy*6C06 digests containing larger quantities of GlcNAc and other oligomers (Fig 2A). Consistently, *Pha*B3M02 and *Alt*A3R04 did not grow on chitobiose as the sole carbon source (Fig 2B, Fig S3). Inspired by this observation, we tested whether the differences in hydrolysis product profiles among degraders could control which exploiters grow in coculture. For this purpose, we grew degraders *Vib*1A01 and *Psy*6C06 in coculture with the four exploiters. In accord with the enzyme digest data (Fig 2A), *Vib*1A01, which produces a range of oligomers supported growth of all species in coculture (Fig 2D). The chitobiose producer *Psy*6C06 was able to support all exploiters except *Pha*B3M02, showing that the composition of degradation products can influence the abundance of specific exploiters. Surprisingly, the exploiter *Alt*A3R04 also grew in coculture with *Psy*6C06, although it grew neither on the enzyme digest of *Psy*6C06 nor on chitobiose (Fig 2b). This suggests that additional metabolites were secreted by *Psy*6C06 or another exploiter to support growth of *Alt*A3R04 but not of *Pha*B3M02, and more generally, that multiple resource pools are formed during chitin degradation that allow downstream growth of less specialized species.

**Figure 2:**
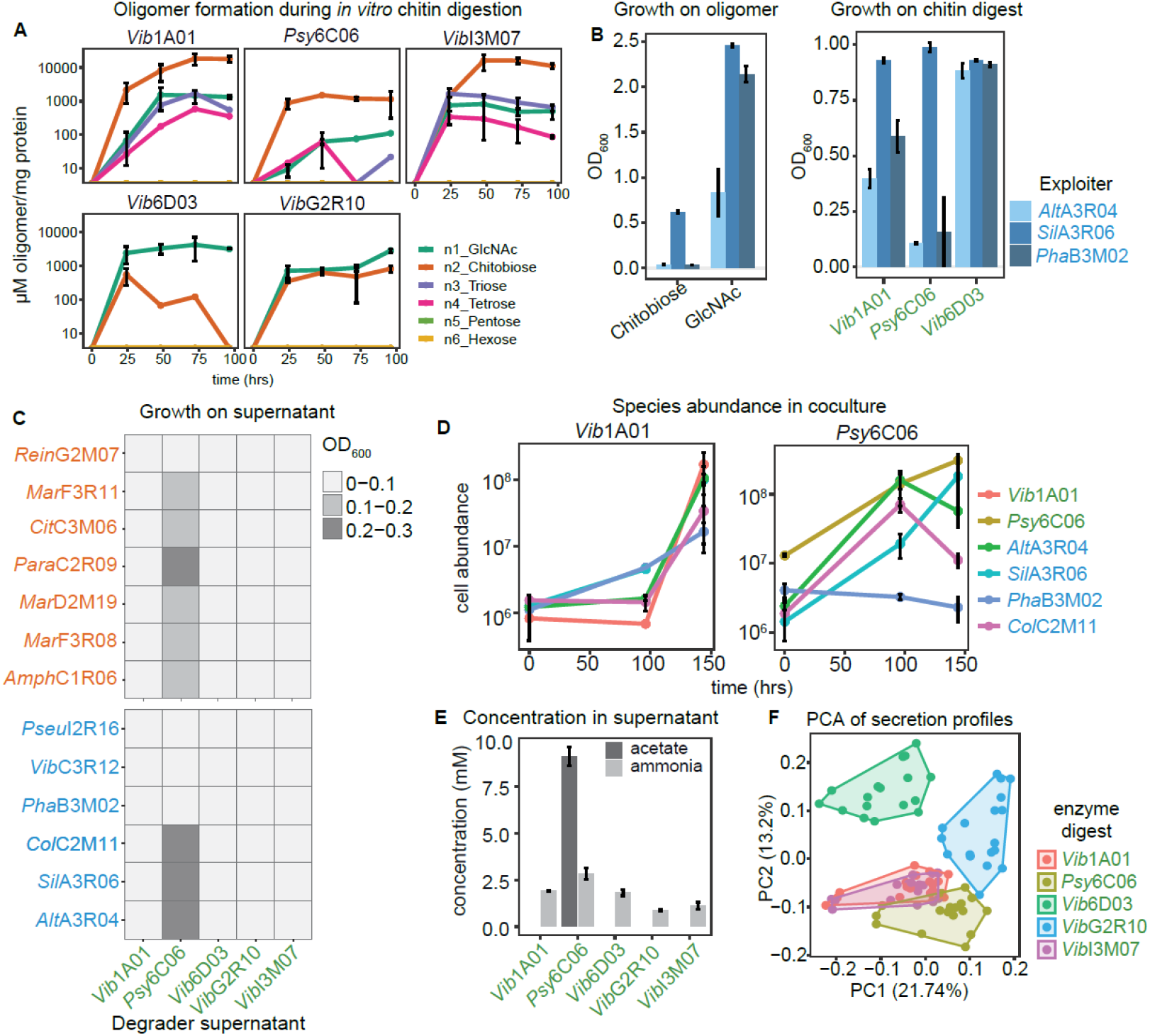
Chitin degradation products influence exploiter population and their secreted resources. A) Quantification of GlcNAc oligonucleotide during *in vitro* digestion of 5 g/l chitin by secreted enzymes from each degrader. B) OD_600_ of three exploiters after 36 h growth on 20 mM GlcNAc, 10 mM chitobiose, and three enzyme digests. C) Growth of scavengers and exploiters after 36 hours on cell-free supernatant produced by each of the five degraders upon growth on colloidal chitin. D) Absolute abundance of species in cocultures of one degrader (*Vib*1A01 or *Psy*6C06) and four exploiters during grown on colloidal chitin, as determined by qPCR. E) Enzymatic quantification of ammonia and acetate in degrader supernatants upon growth on colloidal chitin. F) Principal component analysis exploiter metabolite secretion profiles after growth on each colloidal chitin digest. Colors represent the digest used as a growth substrate. All error bars represent standard deviation from three biological replicates.

To understand the mechanism that facilitated the growth of AltA3R06 in coculture, we investigated whether degraders secrete additional metabolites during growth on chitin to provide a carbon or nitrogen source independent from GlcNAc oligomers. We grew the five degraders on colloidal chitin until early stationary phase and inoculated the cell-free culture supernatants individually with six exploiters and seven scavengers (Fig 2C). While four of the supernatants did not support growth of any species to detectable optical densities, presumably because degraders consumed available limiting nutrients required for growth, *Psy*6C06 supernatant supported the growth of six scavengers and three exploiters. LC-MS targeted metabolomics revealed that GlcNAc and chitobiose concentrations were below the limit of detection in all five supernatants. However, nine metabolites were present in at least one supernatant, including succinate, glutamate, aspartate, and propionic acid (Fig S4, supplemental dataset 1). Since catabolism of GlcNAc into the glycolytic intermediate glucose 6-phosphate entails deacetylation and deamination, acetate or ammonia were quantified with enzyme assays (Fig 2E). Ammonia was likely the nitrogen source provided to other species since it was present at substantial concentrations in all supernatants. Acetate was present only in the *Psy*6C06 supernatant at a concentration 10.9 fold higher than the cumulative molar concentrations of all other detected metabolites. Hence, acetate was presumably the major carbon source in *Psy*6C06 cocultures that fueled growth of *Alt*A3R04 (Fig 2D 2C). Collectively, these results demonstrate that non-degrading community members are selectively supported by a combination of freely available chitin monomers, oligomers and metabolic byproducts secreted during growth of degraders.

We suspected that exploiters also secrete metabolites that less specialized species consume, and wondered if the chitin degradation products influence these secretion profiles. To test this hypothesis, we cultivated each exploiter on colloidal chitin digested with extracellular enzymes obtained from each of the five degraders. After 36 hours, culture supernatants were analyzed with the discovery metabolomics method flow-injection time-of-flight (FIA-QTOF) MS(*38*). Overall, we detected 2521 ions, of which 216 could be annotated based on accurate mass. Principle component analysis using the 216 annotated ions showed a clear clustering of the secretion profiles based on the oligomer substrate (Fig 2F). For example, degraders *Vib*1A01 and *Vib*I3M07 produced similar sets of n1-n4 GlcNAc oligomers (Fig 2A). The secretion profiles of exploiters that used these substrates also showed no clear separation, suggesting that *Vib*1A01 and *Vib*I3M07 exert similar effects on exploiters *via* their similar chitinases. Alternatively, degraders *Psy*6C06 and *Vib*6D03 produced primarily chitobiose and GlcNAc, respectively, and secretion profiles of exploiters grown on these digests clustered distinctly. These results demonstrate that the different chitin degradation products play a major role in determining which metabolites exploiters secrete into a shared resource pool.

### Identification of strain-specific metabolic secretion and uptake profiles

We designed two experiments to identify which of the many secreted metabolites could potentially serve as primary nutrient sources for scavengers. First, all scavengers were cultivated on cell-free supernatants collected from each of the degraders and exploiters after growth on GlcNAc that was used by all species. All supernatants except *Sil*A3R06 supported growth of at least two, but typically more scavengers (5.27±2.4 with OD_600_ > 0.1, Fig 3A), demonstrating secretion of large quantities of metabolites by most species. Measuring 24 absolute metabolite concentrations by targeted LC-MS and enzymatic assays confirmed acetate and ammonia as the major carbon and nitrogen sources, respectively (Fig 3BC, supplemental dataset 1), akin to the results obtained with the *Psy*6C06 chitin supernatant. Yet, additional metabolic niches are manifested by a variety of organic acids, amino acids, and nucleosides that made up as much 31% of the total secreted carbon pool, as in the case of *Col*C2M11. The second experiment was designed to test how these additional metabolites are partitioned among all community members. For this purpose, supernatants of degraders grown on chitin and degraders and exploiters grown on GlcNAc were pooled into a single mixture containing all secreted metabolites. All 18 strains were inoculated in this supernatant and grew individually to densities between 0.15 and 0.7 OD_600_ units, corresponding to 4-6 doublings (Fig S5). To identify consumed metabolites that support this growth, we determined relative changes by FIA-QTOF-MS and searched for metabolites that were i) produced (log_2_ fold change of ion intensity > 1) by degraders or exploiters after growth to late exponential phase in at least one of the cultures, and ii) consumed (log_2_ < −0.5) by at least one species in samples taken over the course of growth in the pooled supernatant (Fig 3D, (Supplementary Dataset 2). Each species had a distinct metabolite consumption profile and a unique ordered hierarchy of which metabolites are consumed first (Fig 3D). This was exemplified by acetate, GlcNAc, pyruvate and glutamate (Fig 3E, Fig S6). For example, *Alt*A3R04 co-metabolized all four substrates, while *Sil*A3R06 and *Pha*B3M02 metabolized them with specific ordered preferences or not at all. These results suggest that species may be specialized to consume unique sets of resources with a distinct hierarchy of preferences, thus enhancing the potential for many species to coexist in environments with fluctuating nutrient availability.

**Figure 3:**
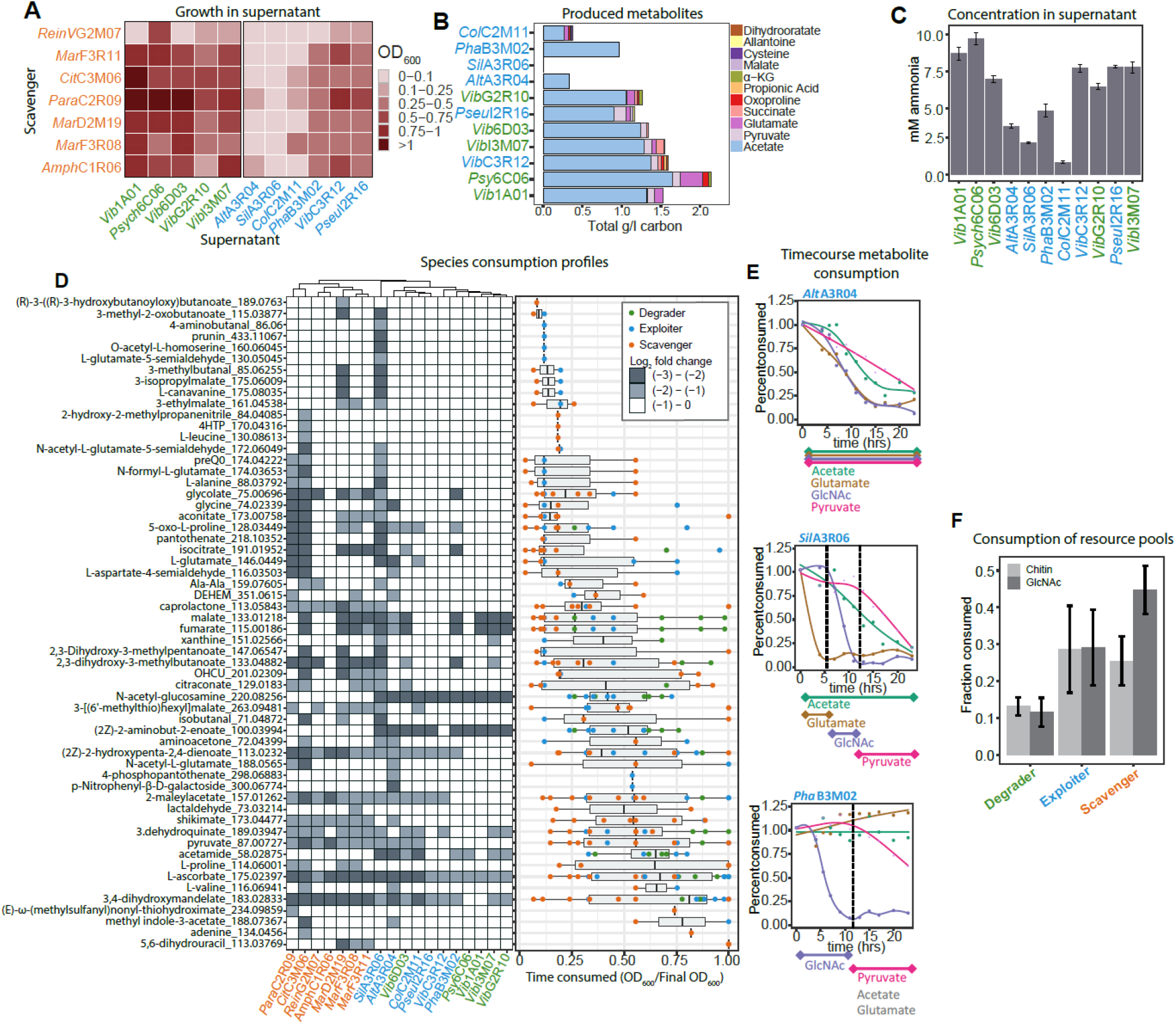
Metabolite consumption and production profiles among community members. A) Growth of scavengers at 36 hours on cell-free supernatants obtained from degraders and exploiters after growth on GlcNAc. B) Metabolite concentration in cultures of degraders or exploiters after growth on 20 mM GlcNAc. C) Ammonia concentration in cultures of degraders or exploiters after growth on GlcNAc. D) Consumption of crossfed metabolites in pooled supernatant. (left) FIA-QTOF-MS measurements of metabolites that are consumed by degraders, exploiters or scavengers (Ion intensity Log_2_ < −0.5). (right) Time course measurements of metabolite consumption. Points represent the fraction of the final OD_600_ at which metabolite consumption is detected for each species E) Relative change of acetate, pyruvate, GlcNAc, and glutamate during growth in pooled supernatant by three exploiters. Curves are fit using local polynomial regression. F) Fraction of available metabolites contained in chitin or GlcNAc derived resource pools that is consumed by each functional guild. All experiments were performed in triplicate. Error bars represent standard deviation. DEHEM, 4-deoxy-α-L-erythro-hex-4-enopyranuronate-β-D-mannuronate; OHCU, 2-oxo-4-hydroxy-4-carboxy-5-ureidoimidazoline. Strain names in green are degraders, blue are exploiters, and orange are scavengers.

Given the sequential colonization of degraders, exploiters, and scavengers (FIG 1C), we wanted to test whether unique metabolic preferences of guilds and species drive this succession. For this purpose, we approximated the availability of resource pools at early timepoints by determining which metabolites are produced by degraders after individual growth on colloidal chitin, since chitin degradation initiates community succession. Conversely, we approximated the resource pools that degraders and exploiters produce at later timepoints by determining metabolites they secreted after growth on GlcNAc. We then calculated the fraction of metabolites that each species consumed from the early or late resource pools (Fig 3F). Of the 32 metabolites present in the chitin-derived resource pool, degraders, exploiters, and scavengers consumed on average 13%, 28%, and 25% of the available metabolites, respectively. Scavengers greatly increased their consumption to 45% of the 54 metabolites in the GlcNAc-derived resource pool, while degrader and exploiter consumption did not change much. Thus, metabolites produced at later timepoints in community succession are consumed preferentially by scavengers, explaining how this functional guild is specialized to thrive on resources that become available at later timepoints in community succession (Fig 1C).

### Metabolic interactions that define outcomes of chitin degrading communities

In light of the unique secretion and uptake profiles of each species, we explored how these traits may contribute to the outcome of communities in coculture. Cocultures were seeded with one of three degraders and one of four exploiters with an additional set of the same four scavengers to create twelve six-member communities (Fig 4A). The three degraders extracellularly produced different sets of chitin degradation products (Fig 2A), which appeared to influence the prevalence of exploiters in several cases. For example, exploiter *Alt*A3R04 used GlcNAc but not chitobiose (Fig 2B) and consistently coexisted well with degrader *Vib*6D03 (Fig. 4A) that produced GlcNAc as a major degradation product (Fig 2A). However, degrader-exploiter coexistence was not entirely explained by their dependence on GlcNAc oligomers. For instance, exploiter *Col*C2M11 did not grow in coculture with degrader *Vib*1A01 (Fig. 4A), despite its ability to grow on chitobiose and GlcNAc formed by *Vib*1A01(Fig 2A, S3). This is likely the result of poor substrate affinity for GlcNAc oligomers and suggests that additional secreted metabolites may fuel *Col*C2M11.

**Figure 4:**
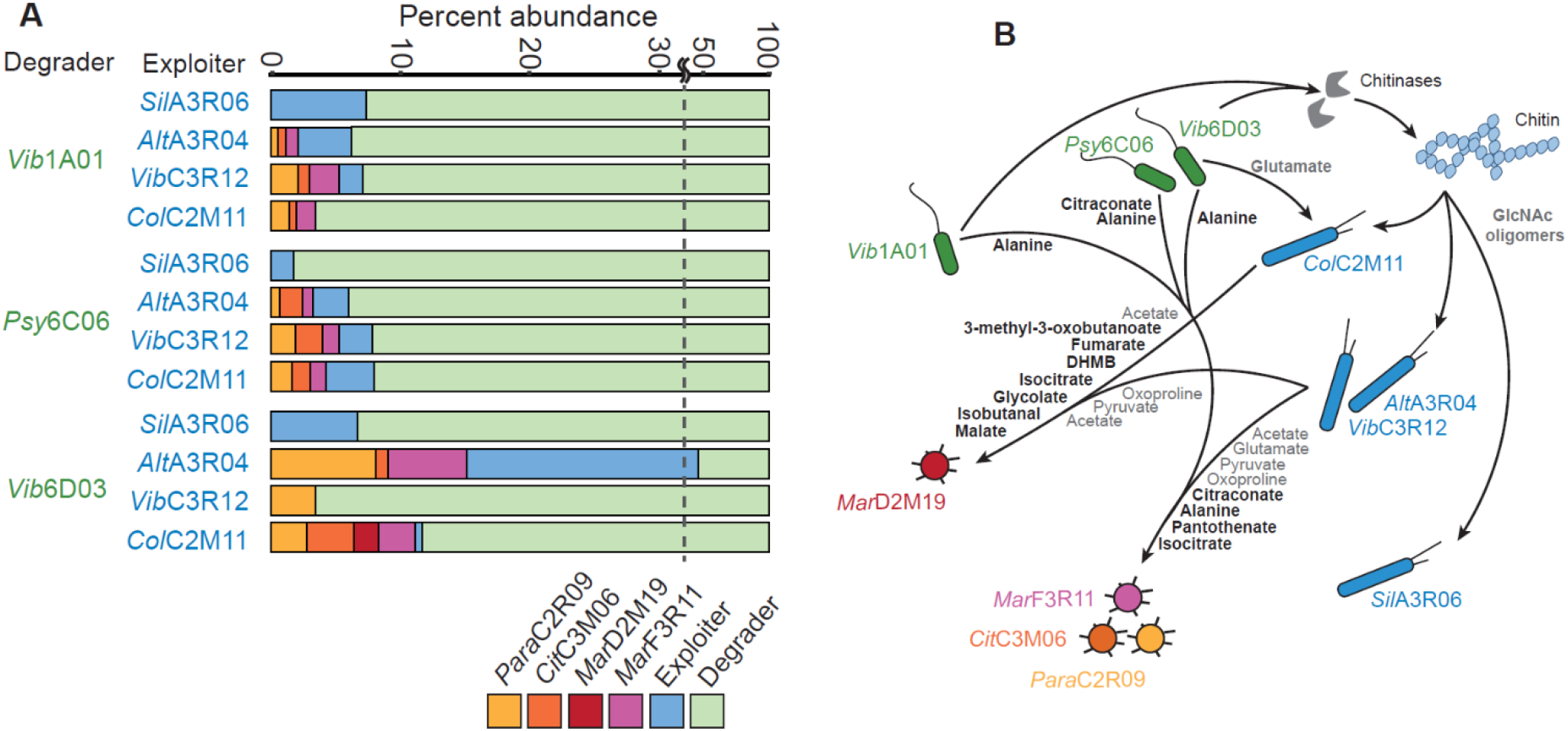
Outcomes of chitin degrading cocultures and underlying metabolic crossfeeding network. A) Each coculture was seeded with six species: one degrader (green), one exploiter (blue), and each of the four scavengers *Para*C2R09 (yellow), *Cit*C3M06 (orange), *Mar*D2M19 (red), *Mar*F3R11 (purple). Cultures were grown for five weeks with four serial dilutions on colloidal chitin and relative abundance is shown for the final culture. Experiments were performed in triplicate. B) Network illustration of metabolites that fuel species growth in the cocultures of panel A. Metabolites commonly secreted by degraders or exploiters and known to be consumed by scavengers are shown in grey. All bolded metabolites were identified using a Boolean modelling approach from the coculture outcomes in panel A. All remaining secreted metabolites and crossfeeding interactions that are not represented in this illustration are contained in (Supplementary Dataset 1, Supplementary Dataset 4). Abbreviation: DHMB, 2,3-dihydroxy-2-methylbutanoate

To gain a comprehensive view of metabolite exchange in cocultures, we compiled metabolite secretion and uptake data of individual species (Supplementary Dataset 3) to construct 12 hypothetical nutrient exchange networks with species combinations based on coculture outcomes observed in figure 4A. Exchanged metabolites had the criteria of being liberated and/or secreted by a degrader during growth on chitin and consumed by an exploiter or scavenger, or being secreted by an exploiter or scavenger and consumed by a scavenger. Of 73 total metabolites included in these networks, each network on average contained 29±8 metabolites and 65±27 exchanges thereof (Supplementary Dataset 4). No scavengers coexisted with *Sil*A3R06 in any coculture (Fig 4A). To assess how exploiters influence the flow of metabolites from degraders to scavengers, we identified potential competition for degrader-derived metabolites by exploiters and scavengers from overlapping consumption profiles (Table S3). On average *Sil*A3R06 consumes 80% of these degrader-derived metabolites, compared to at most 30% by any of the other exploiters. To evaluate whether *Sil*A3R06 itself contributes metabolites to scavengers, we calculated the percentage of metabolites produced by each exploiter that scavengers may consume (Table S4), revealing that *Sil*A3R06 does not secrete any such metabolite. Thus, our hypothetical networks suggest that *Sil*A3R06 likely competes with scavengers for available metabolites without providing any growth-supporting substrates in return. Furthermore, these data illustrate how exploiters have downstream influence on community diversity by changing the resources available to scavengers.

Beyond GlcNAc oligomers and other broadly secreted metabolites such as organic acids that may fuel community succession (Fig 3B), we hypothesized that additional metabolites influenced the survival of species in coculture. To this end, we used a Boolean modelling approach on the hypothetical networks (Supplementary Dataset 4) to identify potential interactions that may explain the presence or absence of species in coculture. We considered metabolites and strains as having only two possible states: present or absent. By comparing metabolite and species state across the 12 networks, we determined which metabolite exchanges were present only in conditions where the presence of a species was observed (Table S2). Figure 4B illustrates the thus inferred flow of metabolites from degraders and exploiters to scavengers in the investigated cocultures (Fig 4A). Among others (Table S2), primarily organic acids, amino acids and one vitamin appear to contribute to community crossfeeding. In one example, glutamate was predicted to feed the exploiter *Col*C2M11 from two degraders but not from *Vib*1A01 that did not coexist with this exploiter, suggesting glutamate to be a growth-supporting metabolite for *Col*C2M11. Also included in figure 4A are metabolites that were commonly produced by degraders and exploiters (Fig 3B) which were shown to be consumed by scavengers (Fig 3D), such as glutamate, pyruvate, oxoproline and the major growth supporting carbon source acetate (Fig 3B). Collectively, these metabolite exchanges illustrate both the potential flow of metabolites that links functional guilds as a whole (Fig 1A, 4B), and how specific metabolites contribute to the presence of individual species.

## Discussion

Resolving microbial community function to the level of contributing species is hampered by the difficulty in monitoring how nutrients are dispersed between species and how individual species can impact the broader community through metabolic interaction networks. Here we demonstrate an approach for how community interaction networks can be learned from the characterization of individual species, and how specialized biopolymer degraders scaffold these networks to shape the population. Degraders secrete chitinases that produce different oligomer profiles that influence the abundance of exploiters, which thrive primarily on oligomers that they cannot generate themselves. Moreover, the composition of oligomer profiles influences which metabolites exploiters secrete into a shared resource pool that is available to the entire population. As such, degraders initiate a hierarchical cascade of nutrient flow into a population.

An open question regarding the exchange of nutrients in a community is the extent to which it is mediated by broad and nonspecific crossfeeding networks(*18*), or by species-specific interactions with an ordered structure(*19*). Our results provide evidence that the exchange of nutrients has an ordered structure that allows individual species to shape the flow of metabolites, and subsequently that nonspecific crossfeeding networks do not capture the complexity of nutrient exchange. We show that a multitude of secreted metabolites provide metabolic niches to support a large population. However, our results further demonstrate that scavengers, which can neither grow on chitin nor its degradation products, preferentially consume metabolites that are formed at later points in community succession, demonstrating how degraders scaffold downstream colonization of non-specialized species. With our approach of integrating individual species consumption and secretion data, we were able to infer metabolic exchange networks that hypothesize specific metabolites that support growth of different species.

Collectively, these results demonstrate how a hierarchical structure is formed within microbial communities by radiating crossfeeding networks. In principle, this crossfeeding structure can be generalized to the multitude of communities that require specialized biopolymer degraders, including those involved in human health(*1*)and biogeochemical transformations(*2*). Although hierarchical structures may be convoluted in more complex environments, we outline a concept for generating hypotheses on crossfeeding networks when constructing smaller consortia that recapitulate key aspect of the entire community.

## Author contributions

S Pontrelli designed and performed all experiments, analyzed data and wrote the manuscript. R Szabo performed genomic analysis of bacterial isolates. S Pollak mapped 16s sequences of bacterial isolates to ESVs of developing chitin communities. J Schwartzman performed 16s amplicon sequencing analysis. D Ledezma performed network analysis of chitin coculture outcomes. O Cordero assisted in data interpretation and writing the manuscript U Sauer designed the experiments and wrote the manuscript.

## Data and materials availability

Metabolomics spectral files can be accessed from the MassIVE database (ftp://massive.ucsd.edu/MSV000087455/ password reviewer123) and will be made publicly available upon publication. The short read 16 s rRNA amplicon sequencing data (BioSample: SAMN11023523-SAMN11023755) have been deposited in NCBI with the BioProject identifier PRJNA478695. All deposited genome sequences have been deposited in NCBI with BioProject identifier and BioSample IDs specified in table S1.

## Supplementary datasets

Supplementary Dataset 1: LC-MS measurements of metabolites produced by degraders and exploiters after growth on chitin or GlcNAc

Supplementary Dataset 2: FIA-QTOF-MS measurements of metabolites produced by degraders and exploiters after growth on GlcNAc or chitin, measurements of metabolites produced by exploiters after growth on enzyme digest of degraders, and measurements of metabolites produced and consumed by all strains after growth on pooled supernatant

Supplementary Dataset 3: Compiled metabolite production and consumption data of each species that participated in 6 member chitin cocultures. This data was used for coculture specific metabolic network construction. The production and consumption data is compiled from individual metabolite production and consumption data of degraders grown on chitin, GlcNAc oligomers that degraders form extracellularly, secretion profiles of exploiters after growth in pooled supernatant, and metabolites consumed by exploiters and scavengers in pooled supernatant.

Supplementary Dataset 4: 12 Hypothetical metabolic exchanged networks that were constructed based on the outcome of six member chitin degrading cocultures.

Supplementary Dataset 5: Proteins related to chitin degradation and metabolism found to be present in the species used in this work.

