## Supplemental information for "Hierarchical control of microbial community assembly"

### Materials and methods

#### Bacterial speciess and chemicals

All chemicals were purchased from Sigma Aldrich unless otherwise noted. GlcNAc oligonucleotide standards were obtained from Omicron Biochemicals. Media used are either Marine Broth 2216 (Difco #279100, Fisher) or MBL minimal media supplemented with a carbon source. MBL is mixed from several stock components: 4x concentrated seawater salts (NaCl 80g/l, $MgCl_2*6H_2O$ 12g/l, $CaCl_2*2H_2O$ 0.6g/l, KCL 2g/l), 1000x concentrated trace minerals ($FeSO_4*7H_2O$ 2.1g/l, $H_3BO_3$ 30mg/l, $MnCl_2*4H_2O$ 100mg/l, $CoCl_2*6H_2O$ 190mg/l, $NiCl_2*6H_2O$ 24 mg/l, $CuCl_2*2H_2O$ 2mg/l, $ZnSO_4*7H_2O$ 144 mg/l, $Na_2MoO_4*2H_2O$ 36mg/l, $NaVO_3$ 25mg/l, $NaWO_4$*2H2O 25mg/l, $Na_2SeO_3*5H_2O$ 6mg/l), 1000x concentrated vitamins (riboflavin 100mg/l, D-biotin 30mg/l, thiamine hydrochloride 100mg/l, L-ascorbic acid 100mg/l, Ca-d-pantothenate 100 mg/l, folate 100mg/l, nicotinate 100mg/l, 4-aminobenzoic acid 100mg/l, pyridoxine HCl 100mg/l, lipoic acid 100mg/l, NAD 100mg/l, thiamin pyrophosphate 100mg/l, cyanocobalamin 10mg/l), 1mM phosphate dibasic, 1mM sodium sulfate,50mM HEPES pH 8.2, and a carbon source. All stocks are diluted accordingly and the volume is adjusted to reach 1x concentrations of all components.

Species were received from the lab of Otto Cordero from Massachusetts Institute of Technology (table S1). All species were previously collected from polymer degrading communities of natural seawater communities from Canoe Beach, Nahant, MA, USA; 42°25'11.5'' N, 70°54'26.0'' W[1,2]. Species were stored in -80°C in glycerol stocks and streaked onto Marine Broth 2216 plates containing 1.5% agar (BD #214010) prior to use.

#### Crossfeeding with pooled supernatant

Single colonies were transferred into liquid Marine Broth 2216 and grown overnight at 220RPM at 27°C. Cells were centrifuged and washed in MBL containing no carbon source two times. This was then used to inoculate the respective culture to a final $OD_{600}$ of 0.01. All supernatants were harvested from species grown in 125ml shake flasks containing 20ml of the respective media and were shaken at 100RPM at 27°C. Supernatants were collected by centrifugation and further filtered using a 0.2 µM PES Stericup Filter unit (Millipore). Supernatants were stored at -20°C until further use. Pooled supernatants were prepared by pooling equal volumes of each supernatant. This includes supernatants collected from each degrader grown on chitin, and each degrader and each exploiter grown on GlcNAc. The pooled supernatant was diluted 25% with fresh MBL containing no carbon source prior to inoculation. Species were inoculated into 1.8ml of pooled supernatant contained in 96 deep well plates with a 4mm glass bead. Plates were shaken at 220RPM at 27°C. Samples were taken every 2-4 hours after inoculation, and cells were removed by centrifugation. Supernatant was stored at -20°C prior to measurements.

#### Preparation of colloidal chitin

10g of chitin powder (Sigma C7170) was added to 100ml of phosphoric acid (85 wt. %) and incubated at 4°C for 48 hours. Roughly 500ml of fresh $H_2O$ was added to this mixture and shaken until the chitin precipitated. This was then filtered using vacuum filter paper (Macherey-Nagel MN615) and fresh water was used to wash the filtrate three times. Chitin precipitate was then added into 3kDa dialysis tubing cellulose membrane (Sigma) and placed in water to remove residual monomers and oligomers. Fresh water was replaced daily three times. Colloidal chitin was then collected, adjusted to pH 7 with NaOH, and homogenized using a Bosch Silent Mixx Pro blender. The colloidal chitin was autoclaved before use.

#### Assay for acetate quantification

The following master mix was used for assays: 100mM Tris pH 7.4, 5mM $MgCl_2$, 1mM phosphoenolpyruvate, 1mM ATP, 3u/ml pyruvate kinase, 3u/ml acetate kinase. The assay was started by adding 20µl of sample to 200µl of master mix. The assay was monitored by measuring an absorbance at $OD_{230}$, and concentrations were determined by comparing the rate of PEP consumption over time compared to acetate standards.

**Assay for ammonia quantification**

The following master mix was used for assays: 100mM Tris pH 8.3, alpha-ketoglutarate 7mM, NADPH 0.25mM, 0.1u/ml glutamate dehydrogenase. The assay was started by adding 20µl of sample to 200µl of master mix. Consumption of NADPH was measured over time by measuring an absorbance at $OD_{340}$, and concentrations were determined by comparing slopes to an analytical standard of ammonium chloride.

**Concentration of extracellular chitinases and enzyme digests**

Degraders were inoculated into 50ml of 2g/l colloidal chitin MBL after growing in Marine Broth 2216 as a preculture. Cultures were grown until mid-late log phase ($OD_{600}$ ~1), and all remaining chitin and cells were removed by centrifugation at 4°C. Supernatant was filtered using a 0.2 µM PES Stericup Filter unit (Millipore). Enzymes were concentrated from the supernatant 50 fold using a 10 kDa centrifugation protein concentration filter (Amicon Ultra, Millipore). Following this, the enzyme was added to fresh MBL containing 5g/l of chitin and sampled every 24 hours to measure for degradation products. The resulting digest was further sterile filtered and diluted by a factor of 2.5 and used to support growth of exploiters. Protein concentration was determined using the Bradford reagent.

**Chitin coculture experiments**

Individual colonies were inoculated into Marine Broth 2216 from solid medium and grown overnight at 220RPM at 27°C. Cells were then washed twice in MBL with no carbon and further inoculated into MBL with 2g/l chitin at $1x10^6$ CFU/ml, except for degraders, which were inoculated at $5x10^6$ CFU/ml for qPCR experiments and $2x10^7$ CFU/ml for 16s coculture experiments. All coculture experiments used 96 deep well plates containing 1.8ml media and a 4mm glass bead. Samples for quantifying species abundance were taken of the planktonic phase. Removal of colloidal chitin from samples was done by centrifuging the cultures at 500rcf for 30 seconds and the top later of culture from each well was sampled. Samples were stored at -20°C. Cell abundance was measured with qPCR or 16s sequencing.

**Genomic DNA purification**

Genomic DNA purification was done using DNAdvance Kit (Beckman A48705) following the manufacturers protocol with the modification that the input for the kit is 100µL of Lysis Master Mix combined with 100µL of bacterial culture that is diluted to an $OD_{600}$ of less than 0.2.

**Cell absolute quantification with qPCR**

qPCR was performed using GoTaq qPCR Master Mix (Promega) following the manufacturer's protocol. qPCR reactions were prepared with a final volume of 15µL using prepared genomic DNA as a template. Samples were measured and data analyzed using a QuantStudio 3 Real-Time PCR System with ΔΔCT method. Absolute quantification of species was obtained by comparing $C_t$ values with those obtained from standard cultures that have known CFU/ml values. PCR primers were designed to have an efficiency of 90-105% and sequences are listed in table S5. Primers were tested for specificity to the bacterial species used in each coculture.

**16s sequencing**

Measuring relative abundance of species in coculture was performed with amplicon 16s sequencing using the MetaFast protocol (Fasteris, Switzerland) including proprietary MetaFast barcoded primers. Amplicons were prepared by amplifying purified genomic DNA samples with a unique pair of MetaFast barcoded primers with Kapa HIFI DNA polymerase. Further library preparation was performed by Fasteris and samples were sequencing with an Illumina MiSeq platform.

Paired, quality-filtered reads obtained from 16S sequencing were processed using dada2, version 1.16 to determine exact sequence variants (ESVs). Except where noted, the default parameters were used[3]. Briefly, sequences were trimmed to remove adapters and further trimmed from positions 25 to 175, yielding 150 bp sequences with quality score >38. The maximum expected error rate was set to maxEE(2,5) to ensure stringent trimming and better differentiate reads belonging to species with very similar sequences. After learning error rates, samples were pooled to define variants. Species were matched to ESVs using a custom BLAST built from the full length 16S genes of species used in the experiment[4]. The taxonomy of the ESVs was further confirmed by taxonomic identification using DECIPHER. The SILVA SSU r138 database from 2019 was used for DECIPHER analysis[5].

**Genome annotation**

The species used for this analysis were isolated and sequenced as part of two experiments previously performed in the Cordero lab for studying the succession of natural microbial communities from seawater on polysaccharide particles[1,2]. For each genome, protein-coding genes were predicted and translated using Prodigal v2.6.3[6]. Predicted protein sequences were compared to a custom database of profile hidden Markov models (HMMs) of proteins involved in growth on chitin using the *hmmsearch* function of HMMER v3.3 with default parameters[7]. Publicly-available HMMs were downloaded from the Pfam v33.1[8] or TIGRFAM v15.0 databases[9] (see Supplementary dataset 5 for accession numbers). Custom HMMs were made by identifying experimentally-verified proteins of interest[10,11], finding their homologs in the UniProtKB/Swiss-Prot v2020_06 database[12], creating a seed alignment using MAFFT v7 with default parameters[13,14], and building the profile HMMs using the *hmmbuild* function of HMMER with default parameters (see Supplementary dataset 5 for details on each custom HMM). The isolate strain's proteins were annotated based on the *hmmsearch* results if the protein length was at least 100 amino acids, the independent E-value was less than $1\times10^{-9}$, and the domain score was greater than 30. Only most significant annotation was used for each protein sequence. Gene copy numbers were calculated for each genome by tallying the number of annotations made for each protein group. Genome sequences for each organism can be found using NCBI accession numbers listed in table S1.

**Metagenomic 16S rRNA dynamics**

The 16S rRNA sequence of each genome was determined by first amplifying it using the 27F (5-AGAGTTTGATCMTGGCTCAG-3) + 1492R (5-GGTTACCTTGTTACGACTT-3) universal bacterial primers(19, 20), followed by sanger sequencing. We mapped the 16S sequences of our isolates back to the metagenomic 16S trajectories (PRJNA319196, PRJNA478695) using blast[4]. For each isolate 16S matches in the metagenome were found using a moving threshold approach – we searched for matches starting at 100% identity and going down to 97% identity at 1% identity steps, and if a match was found in any identity threshold the search was ended and all metagnomic hits above that threshold were considered to belong to that isolate 16S. The read counts were transformed into relative frequencies and only entries with a relative abundance >$10^{-3}$ were retained for downstream analysis. Each experiment was originally performed in triplicate, and for each isolate we take the mean relative abundance at each time-point. For each isolate the trajectories were normalized by their highest relative abundance (so that the highest point of any trajectory is 1), and the average of these normalized trajectories is what is reported in Fig 1C.

**Untargeted metabolite profiling and analysis**

Untargeted methods were carried out using Flow Injection Analysis Quadroupole Time of Flight Mass Spectrometry (FIA-QTOF-MS). All supernatants were first diluted 100fold into water before measurements. Metabolomics was carried out with a binary LC pump (Agilent Technologies) and a MPS2 Autosampler (Gerstel) coupled to an Agilent 6520 time-of-flight mass spectrometer (Agilent Technologies). This was operated in negative mode, at 2Ghz for extended dynamic range, with a m/z (mass over charge ratio) range of 50-1000, as described previously[15]. The mobile phase consisted of isopropanol:water (60:40, v/v) with 5mM ammonium fluoride buffer at pH 9 and the flow rate was 150 µl/minute. Raw data for all measurements was subjected to a spectral processing and alignment pipeline using Matlab (The Mathworks, Natick) as described previously[15].

Overall, we detected 2521 ions, of which 216 could be annotated based on accurate mass, corresponding to a maximum of 526 unique metabolites. Ions were annotated with a tolerance of 0.005 Da against a curated compound library that contains all metabolites predicted to be present in at least one of the species used in this study based on BioCyc databases. If a single m/z ion is matched with multiple isomeric or isobaric compounds within the compound library, the compound that participates in the largest number of enzymatic reactions in pathway databases for all 18 organisms used in this work is chosen as the top annotation. Certain isomers were resolved using LC-MS methods as described below, in which case the correct annotations were manually edited.

**LC-MS exometabolomic compound measurements**

Extracellular metabolomics measurements were performed as described previously[16]. Briefly, chromatographic separation was performed using a Poroshell 120 EC-CN 2.1 x 150 mm, 2.7 µm column (Agilent) in an Agilent 1290 infinity stack set to 40°C. The method performed was isocratic, with a mobile phase consisting of 10mM ammonia acetate pH 5.9 and 5% acetonitrile with a flow rate of 250µL/min and a 5 µL injection volume. An Agilent 6520 series quadrupole time of flight mass spectrometer was used for measurements with a m/z range of 20-400 operated in negative mode. Prior to LC-MS measurements, samples were diluted both 20 and 100 fold in water to maximize the number of measurable compounds within the limits of quantification. Data analysis was performed using Agilent Masshunter Quantitative Analysis software. Peaks were integrated using spectral summation integration with integration windows that are defined based on retention times observed from analytical standards. The Agilent software was used to determine polynomial calibration curves with a goodness of fit ($R^2$ > 0.97).

**GlcNAC oligomer quantification**

GlcNAc oligomers were quantified using LC-MS with an Agilent 1100 series stack. Chromatographic separation used a Poroshell 120 HILIC-Z 2.1 x 100 mm, 2.7 µm column (Agilent) kept at a temperature of 30°C and a constant flow rate of 1ml/min. 5 µL of sample was injected after five-fold dilution in 80% acetonitrile. Mobile phase A consists of 10mM ammonia acetate pH 9 in water, and mobile phase B consists of 10mM ammonia acetate pH 9 in 90% acetonitrile. At injection, the mobile phase contains 90% B phase for 2 minutes, followed by a gradient to 40% B phase until 12 minutes and held for one more minute before being re-equilibrated at the starting condition. An Agilent 6520 series quadrupole time of flight mass spectrometer was used for measurements, operated in negative mode at 2GHz for extended dynamic range, with a m/z range of 100-1500. Calibration curves were measured using analytical standards and used to calculate concentrations of oligomers in the sample.

**Inference of coculture metabolic networks**

Metabolic exchange networks were constructed based on coculture outcomes (Fig 4A) and analyzed with Boolean logic to identify metabolites that contribute to the survival of each species. Networks were constructed using secretion profiles of degraders grown on chitin (Supplementary dataset 1 and 2), GlcNAc oligomers that degraders form extracellularly (Fig 3A), secretion profiles of exploiters after growth on pooled supernatant (Supplementary dataset 2), and metabolites consumed by exploiters and scavengers in pooled supernatant (Fig 3D). Combined metabolite production and consumption data used for construction of metabolic networks is contained in Supplementary dataset 3. The metabolic networks for each coculture used Boolean logic to infer two metabolite pools that encompass the sequential colonization of chitin particles by degraders, exploiters and finally scavengers. In all analysis, metabolites were assumed to follow Boolean logic with only two possible states: present or absent. The first metabolite pool is a degrader-derived metabolite pool consisting of metabolites produced by the specific degrader in the culture and represents resources available to support growth of exploiters. Metabolites that are produced by a degrader (Supplementary dataset 3) are considered to be present in the degrader-derived metabolite pool. The second resource pool is an exploiter-derived metabolite pool that represent metabolites available to support scavengers. Metabolites that are produced by a degrader and consumed by an exploiter (Supplementary dataset 3) are absent, metabolites that are produced by a degrader and not consumed by an exploiter are present, and metabolites that are produced by an exploiter are present.

For exploiters and scavengers that selectively grew in one coculture and not another coculture, Boolean logic was used to identify metabolites that contribute to this selective growth. For exploiters, metabolites that the exploiter can consume (Supplementary dataset 3) were identified that are present in a degrader derived resource pool when the exploiter grew and are absent when an exploiter showed no growth. For scavengers, metabolites were identified which the scavenger consumes (Supplementary dataset 3) that are present in an exploiter derived resource pool when the scavenger grew and absent in a coculture when the scavenger showed no growth. These metabolites are shown in table S2.

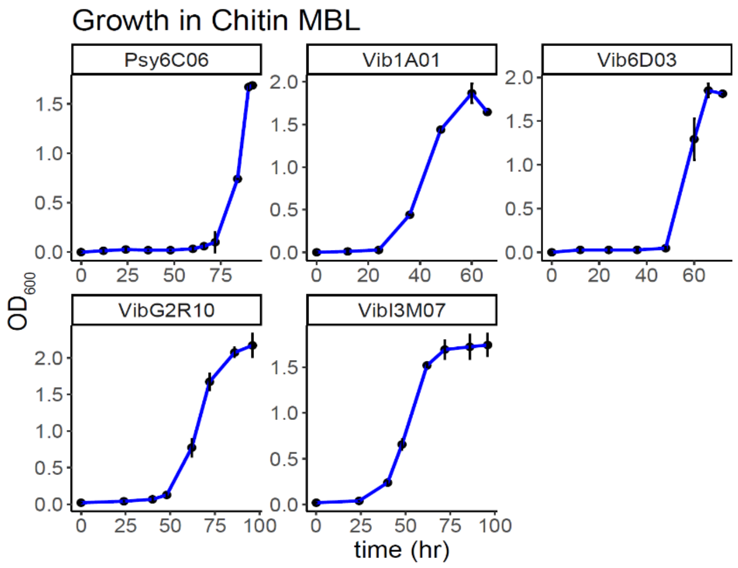


**Fig S1:** Growth of degraders on Chitin MBL containing 2g/l colloidal chitin. Error bars represent standard deviation of three biological replicates

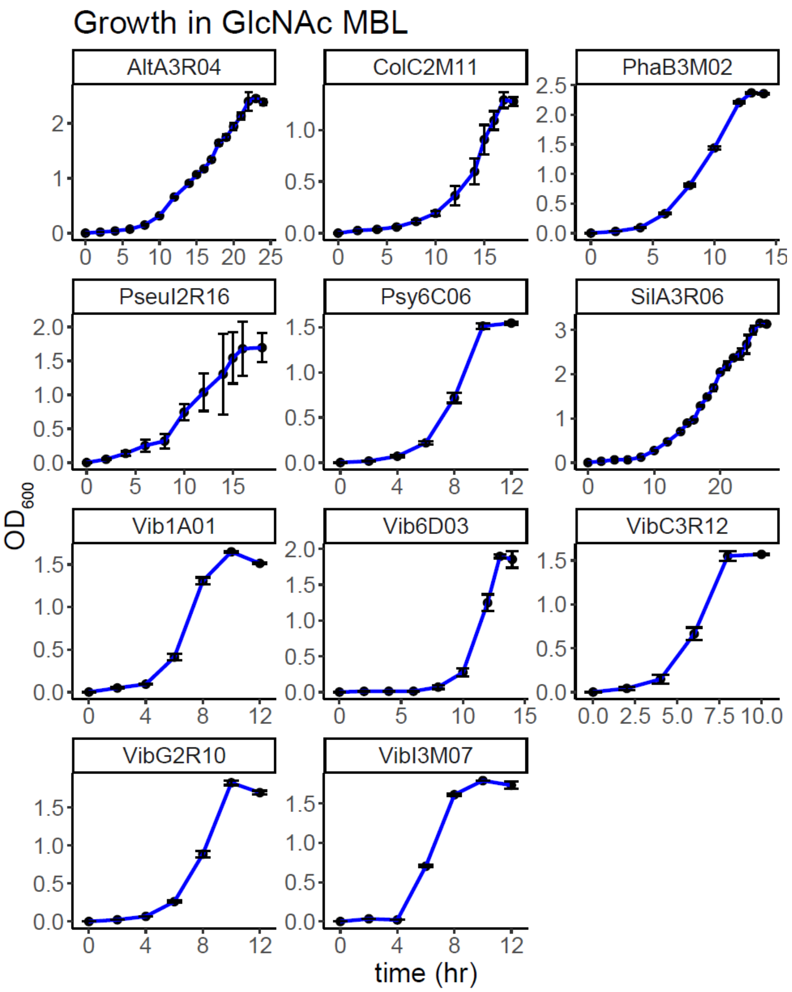


**Fig S2:** Growth of degraders and exploiters on GlcNAc MBL containing 20mM GlcNAc. Error bars represent standard deviation of three biological replicates

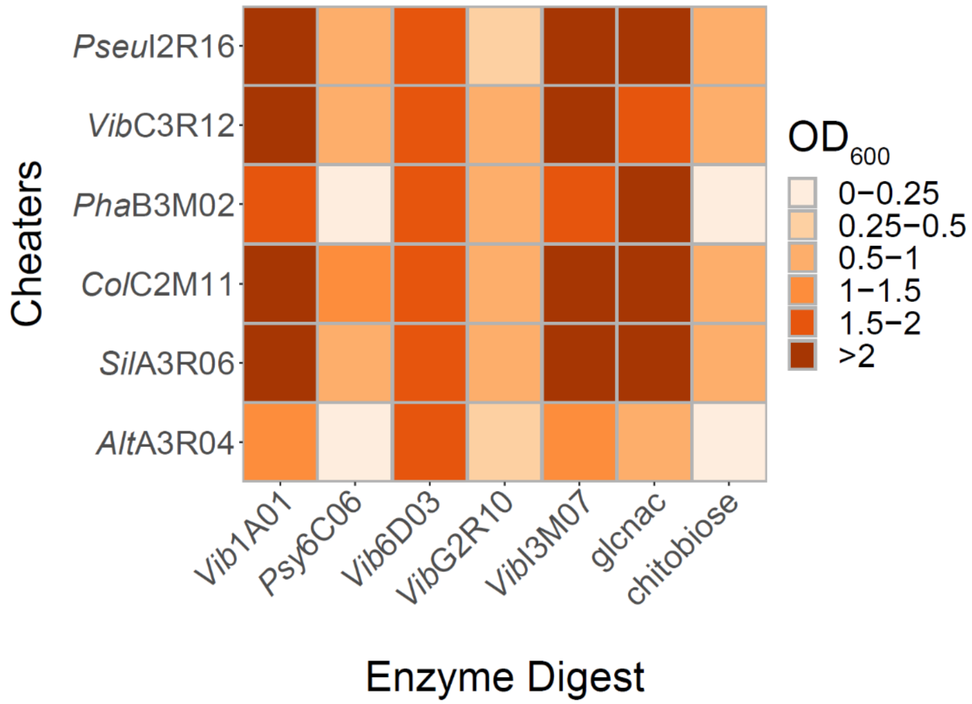


**Fig S3:** Growth of exploiters (vertical axis) after 36 h on colloidal chitin that was digested by enzymes concentrated from supernatants of the five degraders, 20mM GlcNAc or 10mM chitobiose (horizonal axis). $OD_{600}$ data is normalized by the amount of enzyme collected from each degrader as this affects the amount of oligomer liberated from undigested chitin. These values are (left to right) 97.1, 251, 140, 39.3, and 93.6 µg/ml.

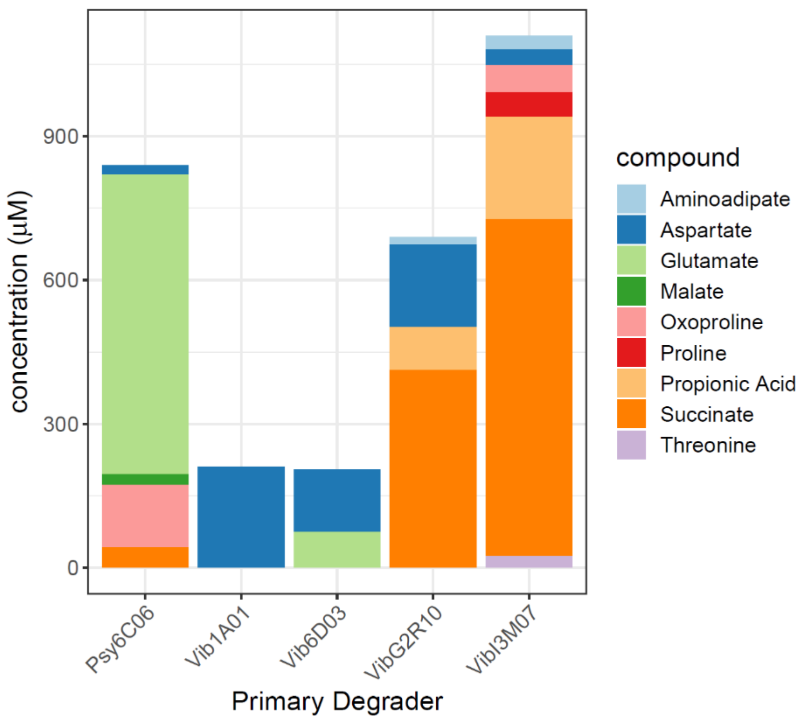


**Fig S4:** LC-MS measurements of metabolites secreted by degraders after growth on Chitin. Measurements represent the average of three independent biological replicates.

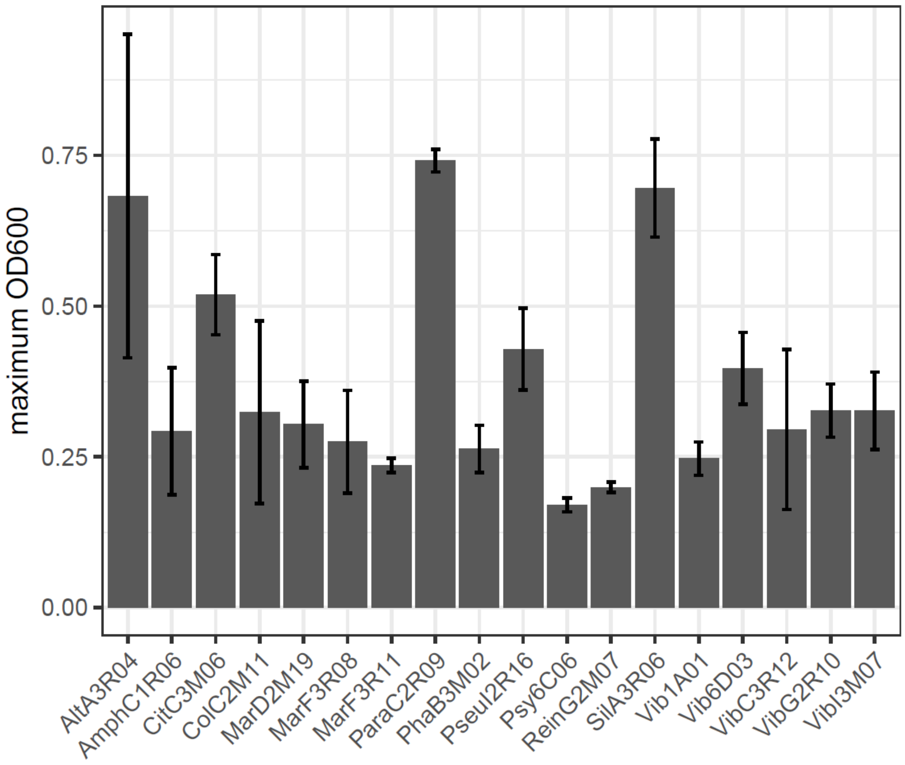


**Fig S5:** $OD_{600}$ of each species after growth on pooled supernatant at 24 hours. Error bars represent standard deviation of three biological replicates.

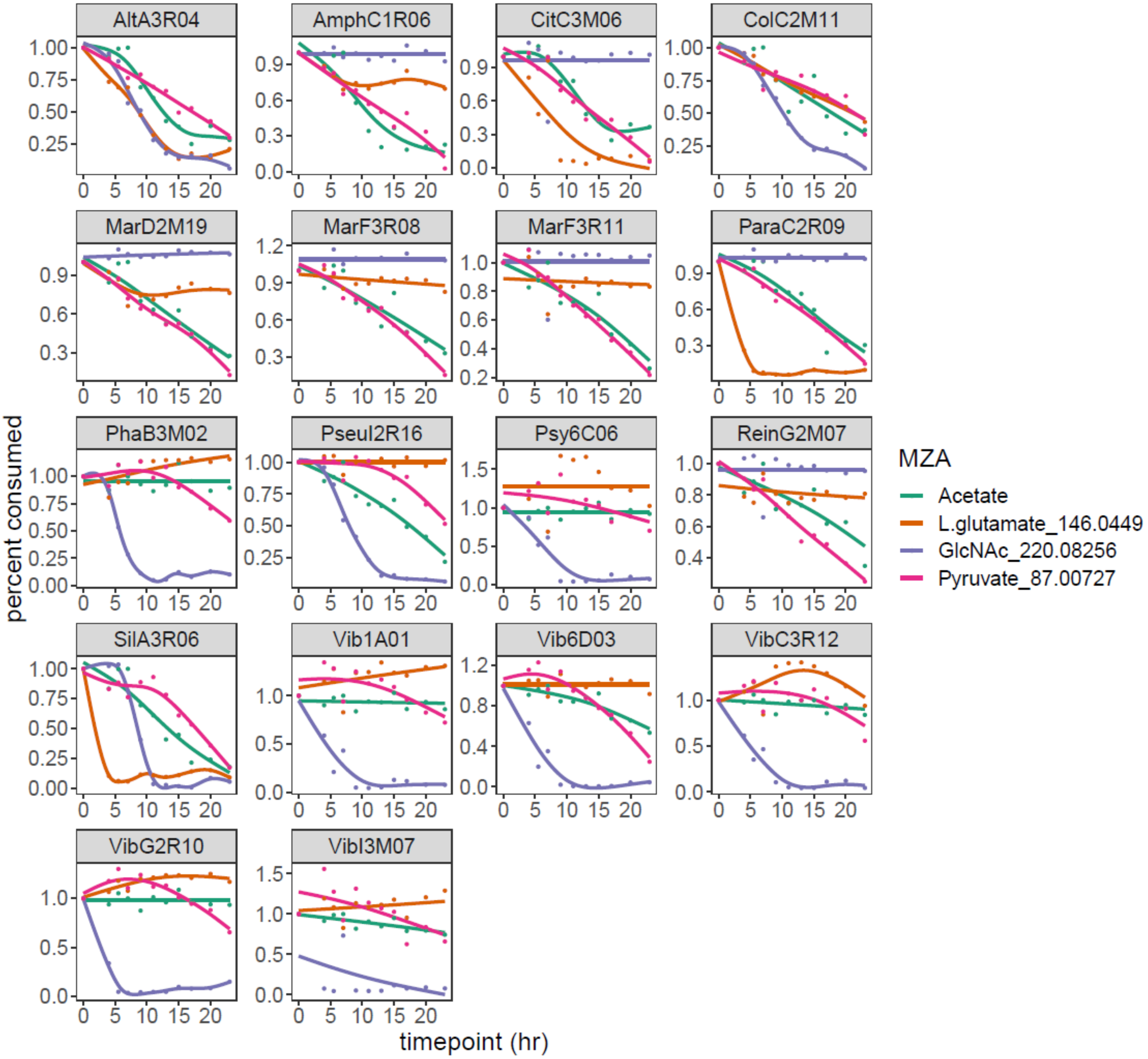


**Fig S6:** FIA-QTOF-MS timecourse measurements of consumption of four metabolites by each species when grown in pooled supernatant. Curves are fit using local polynomial regression.

| Species | Taxonomic ID | Functional Guild | NCBI BioProject | NCBI BioSample |
|---|---|---|---|---|
| AltA3R04 | Alteromonas sp. | Exploiter | PRJNA478695 | SAMN19350919 |
| AmphC1R06 | Amphritea sp. | Scavenger | PRJNA478695 | SAMN19350932 |
| CitC3M06 | Citricella sp. | Scavenger | PRJNA478695 | SAMN19350936 |
| ColC2M11 | Colwellia psychrerythraea | Exploiter | PRJNA478695 | SAMN19350933 |
| MarD2M19 | Marinobacter sp. | Scavenger | PRJNA478695 | SAMN19350941 |
| MarF3R08 | Marinobacter sp. | Scavenger | PRJNA478695 | SAMN19350954 |
| MarF3R11 | Marinobacter sp. | Scavenger | PRJNA478695 | SAMN09522136 |
| ParaC2R09 | Paracoccus kamogawaensis | Scavenger | PRJNA478695 | SAMN19350935 |
| PhaB3M02 | Phaeobacter sp. | Exploiter | PRJNA478695 | SAMN09522130 |
| PseuI2R16 | f__Pseudoalteromonadaceae | Exploiter | PRJNA478695 | SAMN19350960 |
| Psy6C06 | Psychromonas sp. | Degrader | PRJNA414740 | SAMN08130274 |
| ReinG2M07 | Reinekea sp. | Scavenger | PRJNA478695 | SAMN19350955 |
| SilA3R06 | Silicibacter sp. | Exploiter | PRJNA478695 | SAMN19350920 |
| Vib1A01 | Vibrio splendidus | Degrader | PRJNA414740 | SAMN07809270 |
| Vib6D03 | Vibrio penaeicida | Degrader | PRJNA414740 | SAMN08130373 |
| VibC3R12 | Vibrio sp. | Exploiter | PRJNA478695 | SAMN09522132 |
| VibG2R10 | Vibrio sp. | Degrader | PRJNA478695 | SAMN19350956 |
| VibI3M07 | Vibrio sp. | Degrader | PRJNA478695 | SAMN19350961 |

**Table S1:** Species used in this study and NCBI accession numbers for accessing genomic sequences.

| Species | Metabolite | m/z |
|---|---|---|
| *Alt*A3R04 | NA | |
| *Sil*A3R06 | NA | |
| *Col*C2M11 | NA | |
| *Vib*C3R12 | Glutamate | 146.0449 |
| *Para*C2R09 | Isocitrate | 191.01952 |
| | Alanine | 88.03792 |
| | Pantothenate | 218.10352 |
| | citraconate | 129.0183 |
| *Mar*D2M19 | Isocitrate | 191.01952 |
| | 2,3-dihydroxy-3-methylbutanoate | 133.04882 |
| | Malate | 133.01218 |
| | 3-methyl-2-oxobutanoate | 115.03877 |
| | Fumatate | 115.00186 |
| | Glycolate | 75.00696 |
| | Isobutanal | 71.04872 |
| *Cit*C3M06 | Alanine | 88.03792 |
| | Pantothenate | 218.10352 |
| | Citraconate | 129.0183 |
| *Mar*F3R11 | Isocitrate | 191.01952 |
| | Citraconate | 129.0183 |

**Table S2**: Metabolites identified with Boolean logic that are present only in conditions where growth of the noted species is observed.

| | *Sil*A3R06 | *Alt*A3R04 | *Vib*C3R12 | *Col*C2M11 |
|---|---|---|---|---|
| *Vib*1A01 | 86% | 21% | 0% | 0% |
| *Psy*6C06 | 88% | 38% | 0% | 13% |
| *Vib*6D03 | 66% | 33% | 0% | 33% |

**Table S3:** Percentage of metabolites that are consumed by exploiters (top row) from the pool of metabolites produced by degraders that otherwise have the capacity to be consumed by scavengers.

| | A3R06 | A3R04 | C3R12 | C2M11 |
|---|---|---|---|---|
| *Vib*1A01 | 0% | 24% | 31% | 34% |
| *Psy*6C06 | 0% | 22% | 26% | 29% |
| *Vib*6D03 | 0% | 32% | 43% | 47% |

**Table S4.** Percentage of metabolites produced by exploiters that have the capacity to be consumed by scavengers

| Primer target name | Primer sequence |
|---|---|
| 1A01 F | GCACAGACCGTCCAAGAGAA |
| 1A01 R | ACGGTCGCAAGCATATCAGT |
| C1R06 F | GCATTCGTCGTCAGCGTAAC |
| C1R06 R | GCAGGTTTTGCGTCGTAAGG |
| G2M07 F | GCGCTGCACTATAGCCTGTA |
| G2M07 R | CACGGGCTTTGAGGTAGTGT |
| G2R10 F | AGTGAAACCACCAATCGGCA |
| G2R10 R | GCACTATTGCTGTTAGGCGC |
| F3R08 F | AAGACTGCTACAGTGGCCAC |
| F3R08 R | CAGCAACGCCAGAAAAGTCC |
| 6C06 F | AGCACACCCTCGCTCTAAAC |
| 6C06 R | TGAGAAAGTCCTGTTCCGCC |
| A3R04 F | GCTTGCCACCTTTCCCAAAG |
| A3R04 R | TCCGGGCACATTGACAATCA |
| A3R06 F | GAGTTACCCCATGTTCCGCA |
| A3R06 R | AATTGTCATTGCCCGATGCG |
| C2M11 F | TGAAGCGACTTTTGCAGTGC |
| C2M11 R | CTGCAAGGGAGTAATGCGGA |
| B3M02 F | AAAGGCGCTCGTCTTTGGTA |
| B3M02 R | AGATGGGAAAGGTCATGCGG |

**Table S5**: qPCR Primers used in this study
